# Pharaoh ant colonies dynamically regulate colony demography by culling young queen and male-destined larvae

**DOI:** 10.1101/211573

**Authors:** Michael R. Warner, Jessica Lipponen, Timothy A. Linksvayer

**Author notes:** these authors contributed equally to this work (Michael R. Warner, Jessica Lipponen).

## Abstract

The success of social insect colonies is dependent upon efficient and dynamic allocation of resources to alternate queen and worker castes. The developmental and molecular mechanisms regulating the caste fate of individual larvae in response to environmental cues have been the focus of intense study. However, the mechanisms regulating colony-level resource allocation into alternate castes (i.e. caste allocation ratios) are less well studied. Here, we systematically manipulate colony demography to elucidate the social regulatory mechanisms of caste allocation in the ant *Monomorium pharaonis*. We find that differences in caste allocation result from differences in timing and efficiency of culling of very young reproductive-destined larvae, which are always present in colonies. Based on our results, we develop a conceptual model depicting how colonies integrate numerous individual-level caste determination decisions to regulate colony-level caste allocation. We propose that adult workers make decisions about culling larvae based on the ratio of the number of workers to the number of eggs contained in colonies, likely signalled by pheromone present on eggs. This strategy is a bet-hedging strategy which enables the dynamic alteration of colony demography in response to internal and external conditions. The strategy is likely key to the ability of *M. pharaonis* and similar ants to thrive in disturbed habitats and to become widespread invasive species.

**Significance Statement:** The defining feature of social insect societies is the presence of alternate queen (reproductive) and worker (non-reproductive) castes of individuals. The fitness of social insect colonies is dependent upon efficient allocation of resources to alternate castes, particularly in the case of highly polygynous (multi-queen) societies. However, the mechanisms by which such societies regulate caste allocation are largely unknown. In this study, we show that colonies manipulate their production of queens (and also males) versus workers according to the present density of eggs in the colony, which serves as a reliable indicator of queens’ fertility. Provided egg density is high, colonies kill queen-and male-destined larvae; when egg density falls, colonies begin to rear queens and males. This flexible resource allocation strategy is key to the ability of highly polygynous species to thrive in marginal (often human-associated) habitats.

## Introduction

The transition from solitary to eusocial living is one of the major transitions in evolution (Maynard Smith and Szathmary 1997). These evolutionary transitions are characterized by an increase in organism (i.e. colony) size and the specialization of individual components (i.e. division of labor) (Maynard Smith and Szathmary 1997). Further elaborations of eusociality are widespread but are fundamentally rooted in division of labor between highly reproductive (queens, as well as kings in termites) and functionally sterile (worker) castes. (Wilson 1971). The dilemma of how to distribute resources to alternative castes is a decision with important fitness consequences analogous to the allocation of energy to somatic and reproductive tissues in multicellular organisms (Wheeler 1911; Hölldobler and Wilson 2009).

Reproductive caste in most eusocial hymenopteran species is fixed by adulthood (Wilson and Others 1971; Hölldobler and Wilson 1990). An individual’s caste fate results from developmental processes involving the response of innate mechanisms to socially-regulated environmental signals (Wheeler 1986; Linksvayer et al. 2011; Trible and Kronauer 2017; Lillico-Ouachour and Abouheif 2017). These innate mechanisms are hypothesized to include a series of developmental switches. The first switchpoint determines sex by ploidy, as hymenopteran males are haploid and females are diploid (Cook 1993; Cook and Crozier 1995; Beye et al. 2003; Heimpel and de Boer 2008; Klein et al. 2016). Later switchpoints that determine female caste-related characteristics may depend upon larval genotype (Anderson et al. 2008; Schwander et al. 2010) as well as a range of external environmental factors (e.g. nurse-regulated quality and quantity of nutrition (Wheeler 1986; Linksvayer et al. 2011; Trible and Kronauer 2017; Lillico-Ouachour and Abouheif 2017), maternal effects (Schwander et al. 2008; Libbrecht et al. 2013), mechanical signals from nurses (Brian 1973; Suryanarayanan et al. 2011; Jeanne and Suryanarayanan 2011; Penick and Liebig 2012), and temperature (Wheeler 1986; O’Donnell 1998; Libbrecht et al. 2013; Lillico-Ouachour and Abouheif 2017)). Developing larvae may also influence these socially-controlled environmental inputs via begging (Le Conte et al. 1995; Creemers et al. 2003; Kaptein et al. 2005) and pheromonal signaling to nurses (Brian 1975; Slessor et al. 2005; Penick and Liebig 2017).

Societies typically coordinate individual caste determination decisions based on environmental factors to produce emergent colony-level caste allocation differences (Oster and Wilson 1978; Hölldobler and Wilson 1990). Colony-level sex allocation is well studied (Queller and Strassmann 1998; Mehdiabadi et al. 2003) and known to be the result of the interaction between the initial proportion of haploid eggs laid by queens and variable rearing (Bourke 1997; Helms 1999; Helms et al. 2000; Passera et al. 2001; Rosset and Chapuisat 2006; Meunier et al. 2008) or culling of brood by nurses (Aron et al. 1995, 2001; Passera and Aron 1996; Keller et al. 1996; Chapuisat et al. 1997). The best-studied example of colony-level female caste allocation is the regulation of production of alternate worker castes (often called worker subcastes, i.e. minors versus majors) in the ant genus *Pheidole* (Lillico-Ouachour and Abouheif 2017). Worker subcaste ratio in *Pheidole* is heritable (Yang et al. 2004), highly evolvable (McGlynn et al. 2012; Lillico-Ouachour and Abouheif 2017), and responsive to various environmental stimuli (Ito and Higashi 1990; Passera et al. 1996; McGlynn and Owen 2002; Yang et al. 2004). The subcaste fate of individual worker-destined larvae is determined by nutrition (Wheeler and Nijhout 1981) and also depends on the concentration of major-produced pheromones (Wheeler and Frederik Nijhout 1984). As these nutritional and pheromonal inputs are dependent upon colony-level processes and colony demography, the result is colony-level dynamic control of the production of majors versus minors (i.e. "adaptative demography" (Wilson 1985)).

In contrast to sex allocation and worker subcaste allocation in *Pheidole*, the social regulatory mechanisms that link individual-level reproductive (queen versus worker) caste determination to emergent colony-level reproductive caste allocation are largely unknown. Colonies that practice seasonal reproduction typically depend in part on environmental cues for female caste determination, in which larvae (Wheeler 1986) or laying queens (Schwander et al. 2008) must overwinter in order for colonies to produce queens. Alternatively, in societies in which queens cannot form colonies without the aid of workers (i.e. colonies that reproduce by budding), new queens are typically only produced when laying queens are absent or rare (Vargo and Fletcher 1986, 1987; Edwards 1987; Arcila et al. 2002; Brown et al. 2002; Boulay et al. 2009; Schmidt et al. 2011). While the ultimate driver of caste allocation shifts in this case is clear (presence/absence of laying queens), the mechanistic details of these shifts remain unclear. A model that integrates social mechanisms and colony-level control of caste allocation is crucial for understanding the life history of polygynous budding species, for which growth and reproduction are dependent upon numbers of both queens and workers (Pamilo 1991a; Cronin et al. 2013).

Similarly to most polygynous budding ants, colonies of the pharaoh ant *Monomorium pharaonis* only produce new queens (as well as males, who do not disperse (Fowler et al. 1993)) when current queens begin to senesce or die (Edwards 1987; Schmidt et al. 2011). Following queen removal, colonies can rear reproductives (males and new queens) at any time from existing eggs and 1st instar larvae (Edwards 1987; Warner et al. 2016). Studies in *M. pharaonis* have thus far identified roles for eggs (Edwards 1987), laying queens, (Tay et al. 2014; Boonen and Billen 2017), older worker larvae (Warner et al. 2016), and colony size in general (Schmidt et al. 2011) in the regulation of *M. pharaonis* caste allocation, but it is unknown how these factors work together, both with and without queens present.

Here, we disentangle the demographic regulators of caste allocation by systematically experimentally manipulating colony demography. We integrate our findings into a conceptual model and propose that colony caste allocation is dependent upon the ratio between the number of workers and the number of eggs currently contained in the colony. Differences in caste regulation between colonies result from differences in the timing and severity of cessation of culling of reproductive-destined larvae. Altogether, our study begins to elucidate the social regulatory mechanisms by which ant species regulate reproductive caste allocation to respond quickly and optimally to environmental disturbance.

## Methods

### Study species and rearing conditions

We constructed study colonies of the ant *Monomorium pharaonis* by mixing several genetically similar stock colonies originally obtained through the sequential crossing of 8 separate lineages. Colonies were maintained at 27 ± 1 ° C, with 50% RH and a 12:12 LD cycle. Colonies were fed *ad libitum* with an agar-based synthetic diet containing sugars and protein in a 3:1 ratio by mass (Dussutour and Simpson 2008) and supplemented with dried mealworms (*Tenebrio molitor*). Food was provided twice weekly. All surveys and manipulations were performed using dissecting microscopes. We identified larval instar and caste by hair morphology (Berndt and Kremer 1986) and size (Warner et al. 2017). When individual brood stages were manipulated, ants were anesthetized using carbon dioxide and brood was gently moved using paintbrushes. Due to the number of different manipulations we performed, we have included a concise table describing each experiment (Table 1).

**Table 1.** Summary of experiments

| Experiment<br>(number and description) | Demographic Class<br>Manipulated | Colony Production<br>Investigated |
| --- | --- | --- |
| 1. the effect of queen number on colony growth and caste regulation | number of queens<sup>1</sup> | egg number,<br>worker pupae number,<br>caste/reproductive ratios |
| 2a. effect of egg number on caste regulation | number of young brood<sup>2</sup> | caste/reproductive ratios |
| 2b. effect of number of workers on caste regulation | number of workers<sup>3</sup> | caste/reproductive ratios |
| 2c. effect of queen number on caste regulation | number of queens | caste/reproductive ratios |
| 3. culling and caste regulation | queen presence | mortality of young brood |

<sup>1</sup>manipulating the number of queens caused an increase in the number of workers and young brood as well (because colonies grew for six weeks), but we directly manipulated only the number of queens
<sup>2</sup>cumulative number of eggs and 1st instar larvae
<sup>3</sup>we concurrently manipulated the numbers of adult workers, worker pupae, and older worker larvae by varying the volume of total individuals, but have referred to this quantity throughout as “number of workers” for simplicity

### Experiment 1. the effect of queen number on colony growth and caste regulation

First, we aimed to determine the impact of queen number on colony ontogeny and reproductive production. We established 40 total colonies using ~1.2 mL (approx. 175 adult workers) mixed brood and workers and either 1, 2, 4, or 8 queens (10 colonies per queen number). We allowed colonies to grow with queens for 6 weeks and then removed queens to stimulate production of males and gynes (new queens), thereby investigating colony growth as well as reproduction. We counted the number of eggs, adult queens, and worker, male, and gyne pupae weekly. We continued surveys until all pupae had emerged (about 7 weeks after queen removal). For the first two weeks, we corrected the number of queens in colonies for whom the treatment assigned (1, 2, 4, or 8 queens) did not match the number of queens the colony contained.

### Experiment 2. the effect of queens, workers, and eggs on caste allocation

We determined from the first experiment that colonies grown with variable numbers of queens differed in caste allocation when queens were removed (see Results). At the time of queen removal, colonies differed in three aspects: number of workers (approximated by number of worker pupae), number of queens laying eggs, and number of eggs available to rear. We subsequently performed three experiments (labeled 2a, 2b, and 2c) in which we systematically varied one of the three quantities prior to queen removal (described in detail below). We counted gyne, male, and worker pupae number weekly, starting three weeks after queen removal (earliest date of reproductive pupation; personal observation, MRW) until all pupae had emerged (about 7 weeks after queen removal). We surveyed all colonies for numbers of all brood stages and worker adults one day after brood manipulations to verify that colonies differed in only the stated variable of interest.

In each experiment, we manipulated or controlled the cumulative number of eggs and 1st instar larvae (referred to as “young brood”) instead of solely eggs for two reasons: 1) such individuals are typically present together in clumps; breaking apart such clumps caused heavy mortality (JL, personal observation). 2) Colony-level caste regulation occurs during the 1st instar, so eggs and 1st instar larvae are the only individuals in normal queenright colonies that may develop into males and gynes as well as workers (Warner et al. 2016).

### Experiment 2a. effect of egg number on caste regulation

We established 40 colonies with ~1.8 mL (approx. 250 adult workers) mixed brood and workers and 16 queens. Colonies were assigned to pairs containing a donor and a receiver colony. After 2 weeks, we removed ½ the number of young brood from donor colonies and added them to the corresponding receiving colony. We donated either 100 or 300 young brood for each pair. Colonies contained about 400 young brood initially, so we established queenless colonies that contained between 100 and 700 young brood but did not differ systematically in the number of queens who had laid the eggs or in the colony size at time of queen removal. Queens were allowed to lay eggs for 2 weeks (in contrast to 1 week in other experiments) because there were insufficient numbers of young brood for manipulation after one week. While this additional week could affect the number of middle-aged (2nd instar or small 3rd instar) larvae, we previously showed that such larvae do not affect caste allocation (Warner et al. 2016). Furthermore, any differences between colonies would occur independently of the final number of young brood, our variable of interest.

### Experiment 2b. effect of number of workers on caste regulation

We established 10 colonies with ~0.6 mL (approx. 125 adult workers) and 10 colonies with ~1.8 mL (approx. 250 adult workers) of mixed brood and workers. We removed all queens immediately and standardized the number of young brood to 150. Rather than manipulating the quantity of individual classes of worker brood or adults, we aimed to establish colonies with systematically larger numbers of workers of all stages, as would be expected for the colonies grown with more queens for 6 weeks as in experiment 1. While these colonies differed in the number of adult workers, worker pupae, and older worker larvae, we will refer only to the number of workers for simplicity. For reference, “small” colonies contained 127 ± 11 (mean ± SE) adult workers and “large” colonies contained 263 ± 8.

### Experiment 2c. effect of queen number on caste regulation

To investigate the direct effect of queen number on caste allocation, we established 36 colonies with ~1.2 mL (approx. 175 adult workers) mixed brood and workers and either 2 or 8 queens each in two sets: one set of 5 two-queen and 5 eight-queen colonies and a second set of 12 two-queen and 14 eight-queen colonies. In the first set, we removed queens after 7 days and standardized the number of young brood to 150. In the second set, we removed queens after 14 days and standardized the number of young brood to 100 due to a general reduction in egg-laying rate.

### Experiment 3. culling and caste regulation

To further investigate how caste regulation occurs in the presence of queens, we measured mortality of young brood in colonies with and without queens. If female caste allocation is regulated by culling young brood, we would expect increased mortality of such individuals in colonies containing laying queens, under the hypothesis that such colonies cull gyne-destined as well as male-destined 1st instar larvae. Because *M. pharaonis* colonies, like many other ant species, generally exhibit a continuous distribution of brood stages (Peacock and Baxter 1950), measuring mortality of specific stages is difficult. Therefore, we established a protocol (described below) that allowed us to follow a specific age-matched cohort of eggs across development to measure mortality.

We constructed 36 colonies with 60 large 3rd instars, 60 worker pupae, 50 queens, and approximately 300 workers each. After 24 hours, we removed the queens and standardized the egg number to 150. After 7 days, we replaced the queens in 18 colonies and left the other 18 colonies queenless. At this time we also added 60 large 3rd instars to all the colonies, as we have previously shown that the presence of large 3rd instars is important to the production of reproductives (Warner et al. 2016), and the large 3rd instars from the initial setup were now gone (either developed into pupae or deceased). In this way, we created queen-present and queen-absent colonies in which we could monitor mortality of the original 150 eggs laid. We surveyed the colonies for all brood stages at the time of queen replacement, and weekly thereafter. In some cases, there was high mortality of eggs and first instars, and there were eight colonies with fewer than 10 young brood at the time of queen replacement; after these colonies were removed from the study, we proceeded with 28 colonies (14 queen-present and 14 queen-absent).

### Statistical analysis

We performed all statistical analyses using R, version 3.4.0 (R Core Team 2014). We fitted generalized linear models (GLM) for production of pupae, caste ratio, reproductive ratio, or proportion of surviving individuals, using treatments such as worker number (defined as a factor: small or large), queen number, young brood number (defined continuously), and queen presence as predictors where appropriate. We defined caste ratio as gynes/(gynes+workers), and reproductive ratio as (gynes+males)/(gynes+males+workers). Note that while these statistics are actually proportions and not ratios, we have labeled them ratios throughout to be consistent with the vast majority of social insect literature. We analyzed caste ratio and reproductive ratio using binomial distributions and counts of pupae using poisson (or quasipoisson when data were overdispersed) distributions. The significance of models was compared using a likelihood ratio test (LRT). A Tukey’s post hoc Test was implemented using the R package multcomp (Hothorn et al. 2008). All plots were generated using ggplot2 (Wickham 2016). All data and R scripts are included as supplemental material.

## Results

### Experiment 1: the effect of queen number on colony growth and caste regulation

Colonies that produced new reproductives in the presence of laying queens were established with fewer queens on average (2.47 vs 5.10; Wilcoxon rank-sum test; W = 310, P < 0.001; see Fig. S1a). Outside of single-queen colonies, which often became too small to produce reproductives during the study period, reproductive production in the presence of queens was generally linked to number of eggs contained during caste regulation, which occurs during the 1st larval instar, approximately 2-3 weeks prior to reproductive pupation (Fig. S1b). When treatment as well as numbers of egg and worker pupae quantified at 2, 3, and 4 weeks prior to survey date were included as predictors of a colony’s likelihood to contain reproductive pupae, only queen number and egg number 2 weeks prior to the date affected the likelihood of reproductive production (LRT; queen number χ^2^ = 13.084, P < 0.001; egg number two weeks prior χ^2^ = 11.225, P < 0.001; all others P > 0.05); more eggs two weeks prior as well as higher queen numbers reduced the probability of a colony containing reproductives (binomial GLM; queen number z = -3.168, P = 0.002; egg number two weeks prior z = -3.161, P = 0.002, all others P > 0.05). Quantitatively, the caste ratio and reproductive ratio produced by colonies were negatively correlated with the number of eggs contained in the colony two weeks prior (Fig. 1; pearson correlation caste ratio: cor = -0.257, P < 0.001, reproductive ratio: cor = -0.278, P < 0.001). Lastly, these early bouts of reproductive production exhibited much lower caste ratios (i.e. relatively fewer gynes produced) than those produced by colonies after queens were removed completely (LRT χ^2^ = 1701, P < 0.001; binomial GLM z = -37.73, P < 0.001).

**Fig. 1.**
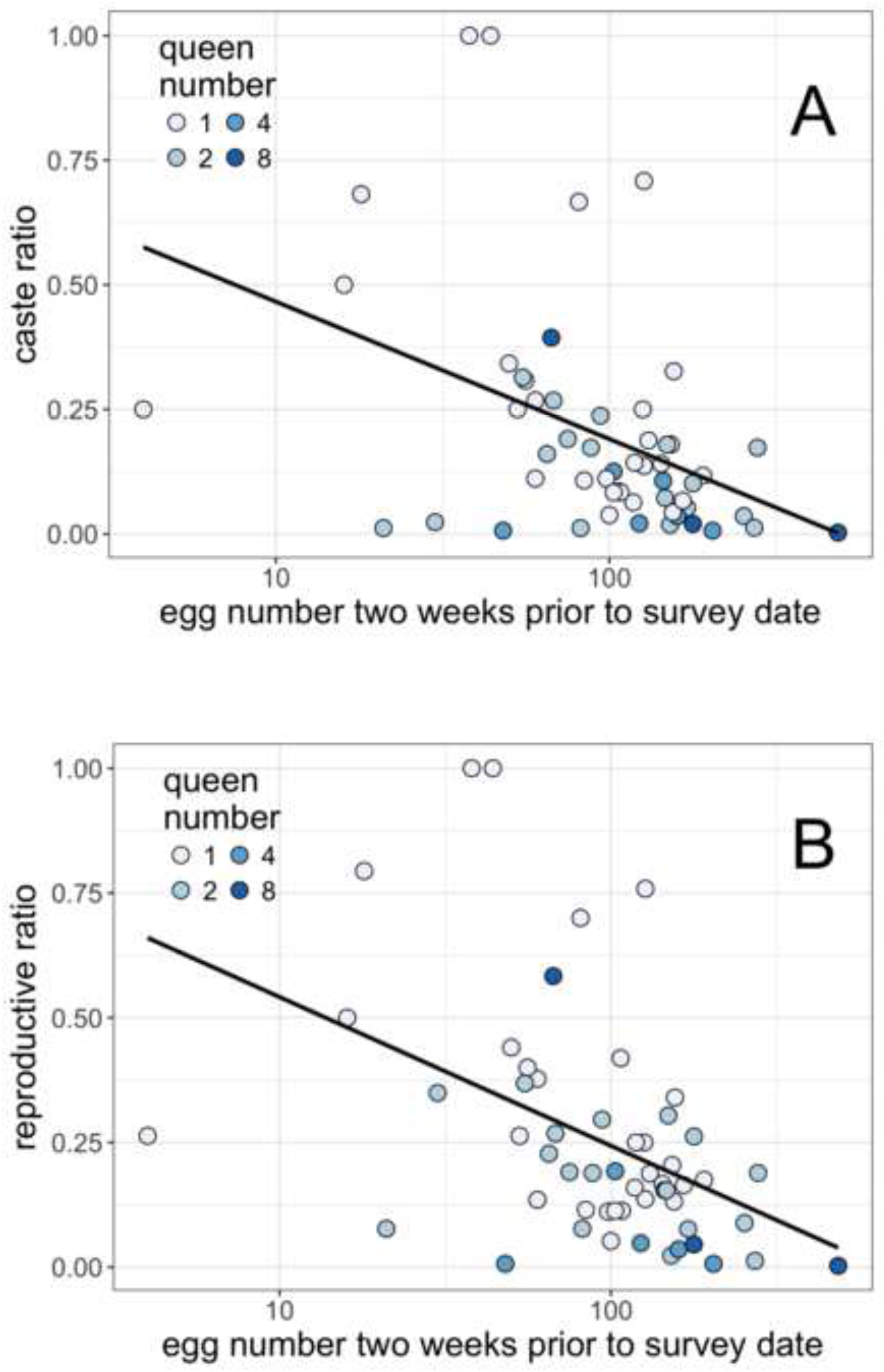
The number of new queens and males produced by colonies containing laying queens (as measured at a particular date) was dependent upon the number of eggs colonies contained two weeks prior to the survey date. Two weeks is approximately the time larvae of the 1st instar (the period of caste regulation) take to pupate, allocation was linked to the number of eggs present at the timing of caste regulation. A) caste ratio (gynes/[gynes+workers]), and B) reproductive ratio ([gynes+males]/[gynes+males+workers]) among all brood produced were negatively correlated to the number of eggs present two weeks prior to survey date (pearson correlation, p < 0.001)

At the time we removed queens from colonies (six weeks after establishment), colonies started with variable numbers of queens differed in size in three ways we measured: number of eggs (Fig S2a; LRT χ^2^ = 39.536, P < 0.001), number of worker pupae (Fig. S2b; LRT χ^2^ = 27.433, P < 0.001), and of course number of queens once we excluded colonies that produced reproductives early from the analysis. Experimental colonies set up with more queens grew more rapidly than colonies with fewer queens, where growth was measured by the number of worker pupae (GLM with quasipoisson link function; t = 6.187, P < 0.001) and by number of eggs present (GLM with quasipoisson link function; t = 5.275, P < 0.001). Colonies differed in reproductive ratio and caste ratio according to number of queens present initially (LRT; reproductive ratio: χ^2^ = 19.552, P < 0.001, caste ratio: χ^2^ = 24.507, P < 0.001). Specifically, colonies started with 2 queens produced higher reproductive ratios (i.e. relatively more reproductives) and caste ratios (i.e. relatively more gynes) than those started with 4 or 8 queens (Fig. S3a). This occurred because colonies with 2 queens produced fewer workers but did not differ in production of reproductives (Fig S3b).

### Experiment 2. the effect of the number of queens, workers, and eggs on caste allocation

In the following series of three experiments, we aimed to disentangle three components of colony size (queen number prior to removal, number of eggs, and number of workers) to evaluate their impacts on caste allocation. Note that in the experiments where we manipulated the total number of workers in colonies, we also varied the number of worker pupae and older worker larvae, so that “number of workers” in these studies refers not just to the number of adult workers, but also including the number of worker pupae and older worker larvae in the colony. Similarly, we refer to the cumulative number of eggs and 1st instar larvae as “young brood”.

Colonies with more young brood but equal worker and queen numbers produced lower caste ratios (Fig. 2a; LRT χ^2^ = 19.185, P < 0.001, GLM z = -4.142, P < 0.001) and lower reproductive ratios in general (Fig. 2b; LRT χ^2^ = 23.247, P < 0.001, GLM z = -4.812, P < 0.001). This effect was likely due to an increase (though non-significant) in worker production with increasing number of young brood (Fig. 2c; pearson correlation r = 0.281, P = 0.079), but not gyne or male production (P > 0.5).

**Fig. 2.**
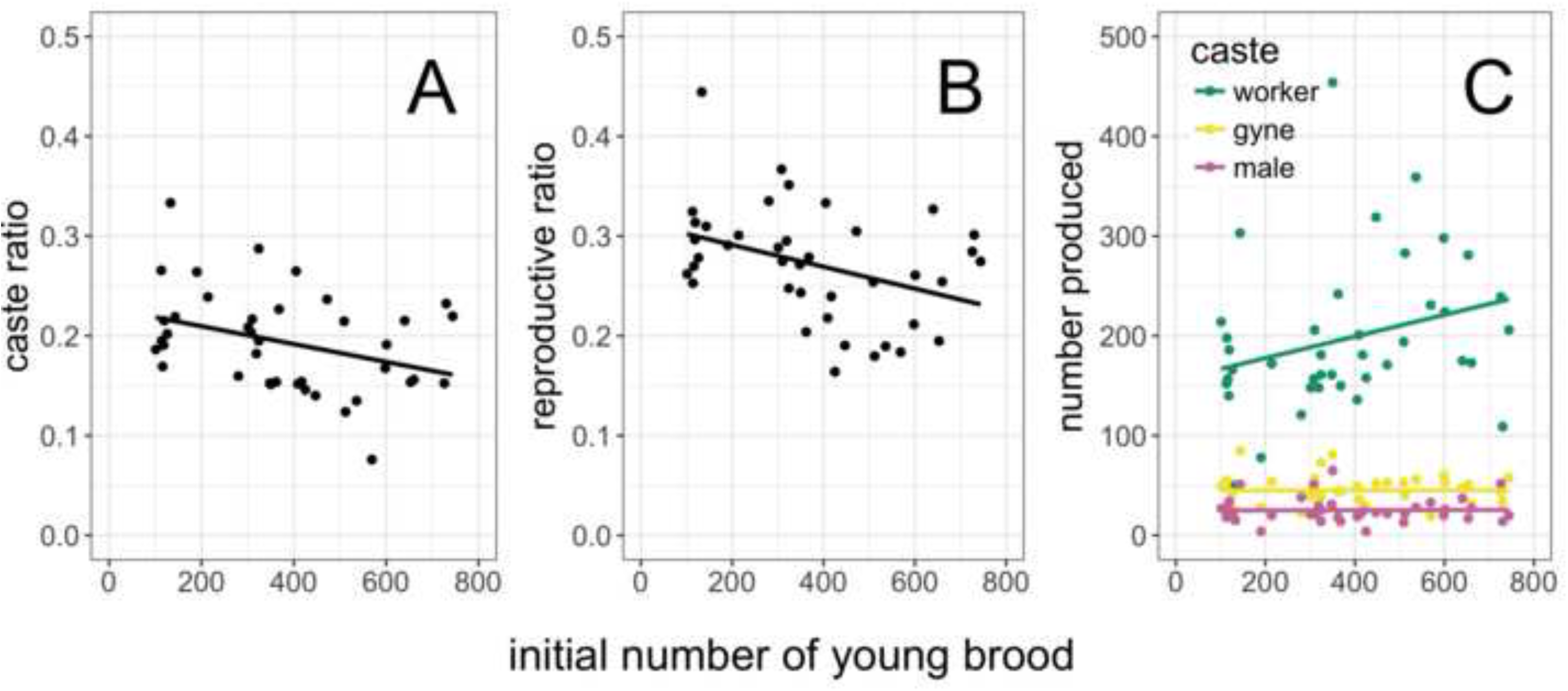
When colonies were established with equal numbers of queens and workers but variable numbers of young brood (eggs and 1st instar larvae), the initial number of young brood affected caste allocation. There was a negative relationship between initial number of young brood and A) caste ratio (gynes/[gynes+workers]) as well as B) reproductive ratio ((gynes+males)/(gynes+males+workers)). This occurred due to a positive relationship between initial number of young brood and worker pupae production but not male or gyne (C)

When colonies were established with equal numbers of queens and young brood, the number of workers affected caste ratio (LRT χ^2^ = 6.408, P = 0.011) and reproductive ratio (LRT χ^2^ = 9.598, P = 0.002). “Small” colonies (containing 127 ± 11 adult workers) produced smaller proportions of each (Fig. 3a). “Large” colonies (containing 263 ± 8 adult workers) produced more individuals of each caste/sex (Fig. 3b; GLM; LRT P < 0.001), but showed a greater increase in gyne and male production (reflected in differing allocation ratios). Queen number itself did not affect the production any caste or caste or reproductive ratio (Fig. S4; GLM; LRT P > 0.5).

**Fig. 3.**
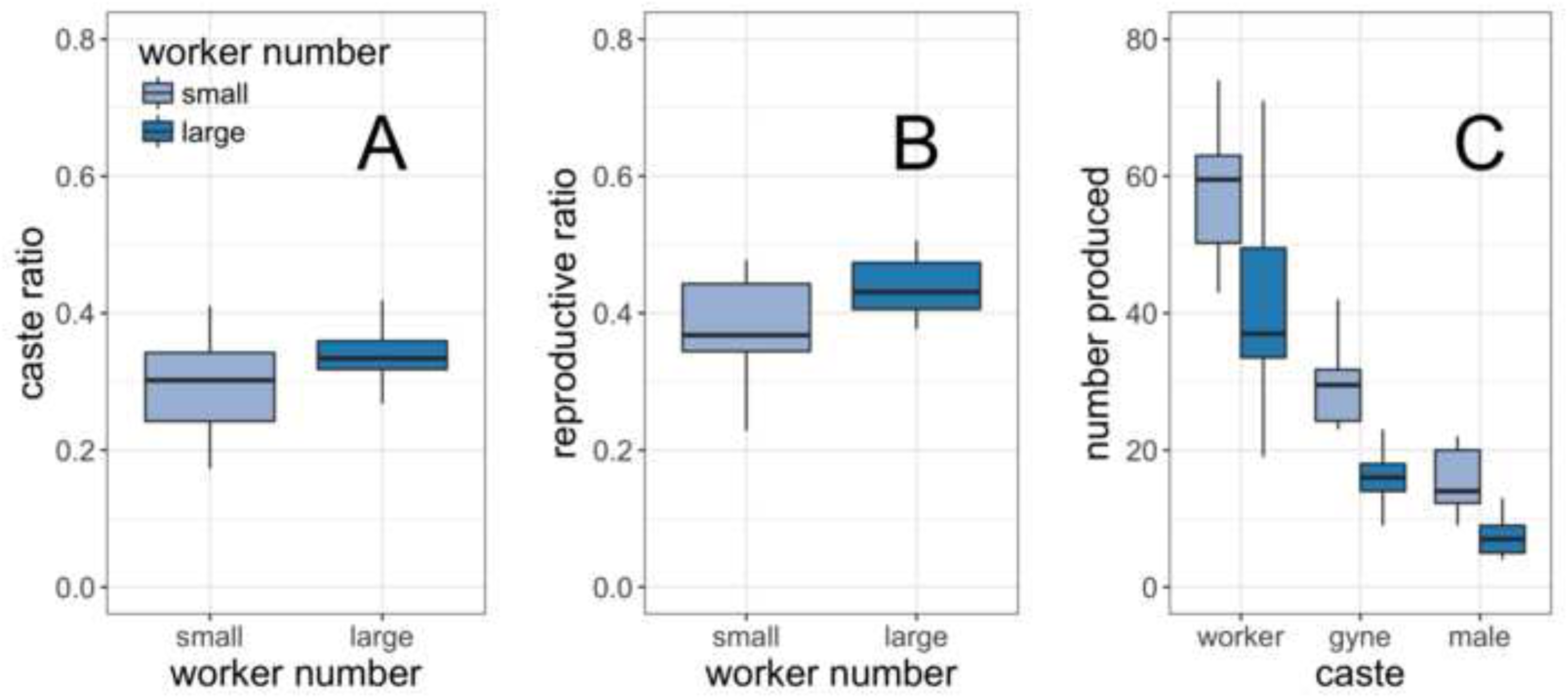
When colonies were established with equal numbers of queens and young brood but variable worker number (small: 127 ± 11 adult workers; large: 263 ± 8), larger colonies produced higher A) caste ratios (gynes/[gynes+workers]) and B) reproductive ratios ((gynes+males)/(gynes+males+workers)). C) While larger colonies increased production of all three castes, the increase was smallest in the case of workers

Colonies of *M. pharaonis* produce reproductives when an inhibition signal, likely a fertility signal on eggs, is reduced (Edwards 1987). To test if timing of the reduction in inhibition affected allocation to reproductives, we compared the average date after queen removal that reproductives pupated (estimated through weekly surveys) to the demographic factors we focused on in the second set of experiments. We reasoned that if the inhibition of reproductive production was halted later, this would be reflected by lower average pupation times. The initial number of young brood (the putative source of the inhibition signal) was positively correlated with timing of male and gyne but not worker pupation (Experiment 2a, Fig. 4a; pearson correlation gyne: r = 0.409, P = 0.009, male: r = 0.319, P = 0.045, worker: r = 0.131, P = 0.420), reflecting the impact of young brood on caste regulation. The number of workers in colonies with equal numbers of young brood impacted the mean timing of gyne but not male or worker pupation; gynes (and males, though the difference was non-significant) pupated later in smaller colonies, i.e. those which had fewer workers per egg (Experiment 2b, Fig. 4b; mann-whitney U-test gyne: U = 88, P = 0.020, male: U = 75.5, P = 0.159, worker: U = 37, P = 0.223).

**Fig. 4.**
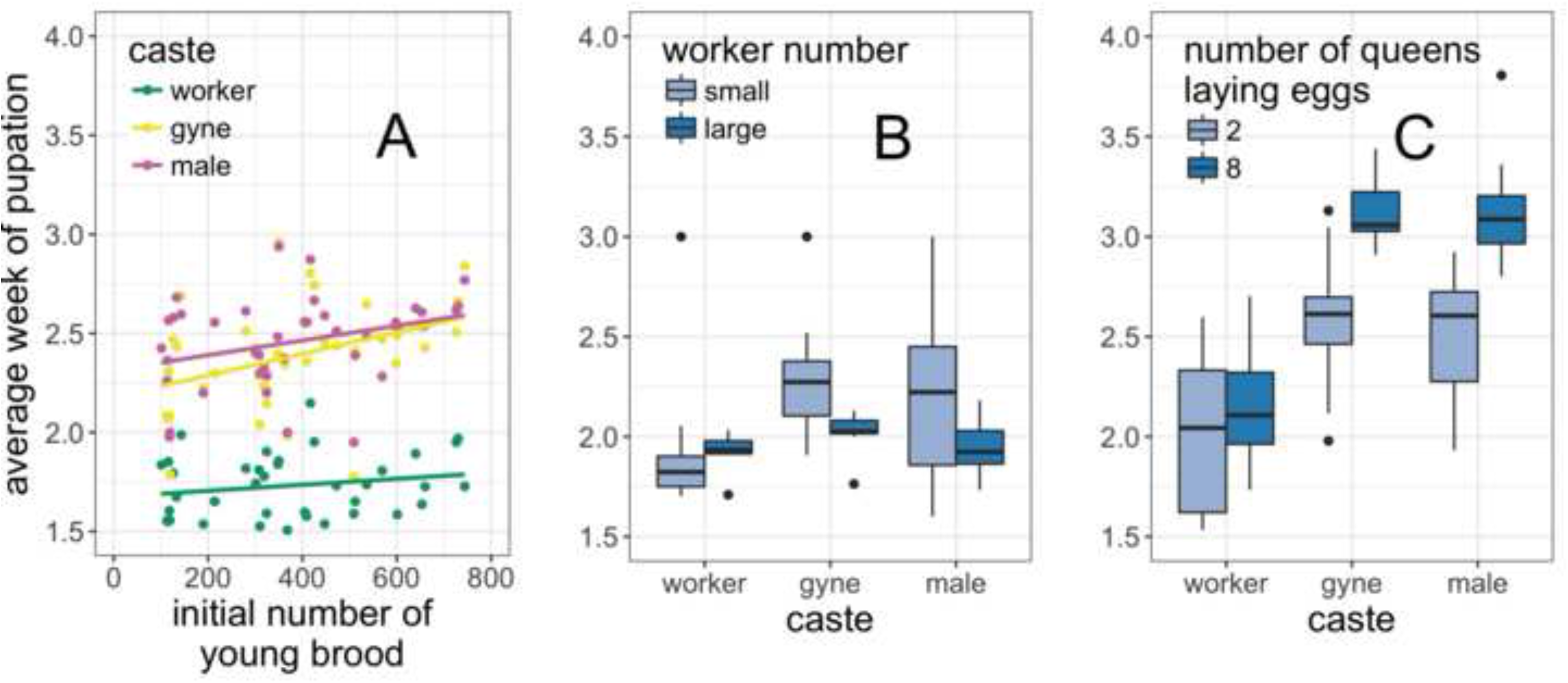
Colony demography affected the average time at which individuals began to pupate (that is, the average number of weeks after queen removal that we observed pupae in experimental colonies). Male and gyne pupae were observed later in colonies containing A) more eggs, B) fewer workers (“small”: 127 ± 11 adult workers; “large”: 263 ± 8), and C) in which eggs were laid by eight versus two queens. Later pupation times for reproductive pupae likely indicate that colonies began rearing reproductive pupae later

Surprisingly, when worker and young brood number were experimentally controlled, the number of queens present in colonies also affected the mean time of pupation of gynes and males but not workers (Experiment 2b, Fig. 4c; mann-whitney U-test gyne: U = 305, P < 0.001, male: U = 312, P < 0.001, worker: U = 197, P < 0.001). While this change in timing seems discordant with our previous analysis of the link between caste regulation and timing of pupation, it is possible the colonies differed nutritionally. While colonies contained similar numbers of eggs and workers, it is likely that being reared with 8 queens reduced resources stored by colony members, such as in replete workers (Børgesen 2000). When colonies with 8 queens started producing reproductives, they may simply have lagged behind because they couldn’t provision larvae as quickly. Rather than cull larvae they could not provision, they simply took longer to rear them, as is witnessed in the argentine ant *Linepithema humile* (Aron et al. 2001). This is consistent with regulation occurring through a cessation of culling driven by demography, rather than by the temporary nutritional state of the colony.

### Experiment 3: culling and caste regulation

Mortality of eggs and 1st instar larvae was higher in colonies in which queens were reintroduced (Fig. 5; GLM; LRT χ^2^ = 41.243, P < 0.001), where survivorship was defined as the proportion young brood that had molted to subsequent larval stages (2nd and small 3rd) one week after queens were replaced. Notably, queen-present colonies produced exclusively workers. Therefore, our data are consistent with increased mortality of eggs and 1st instar larvae in queen-present colonies. It was not possible to definitively compare survival to the adult stage between queen-absent and queen-present colonies because, by the time focal individuals began to pupate, there was a continuous distribution of brood in colonies that contained queens. This made it impossible to further track the survival of a cohort of young brood in queen-present colonies.

**Fig. 5.**
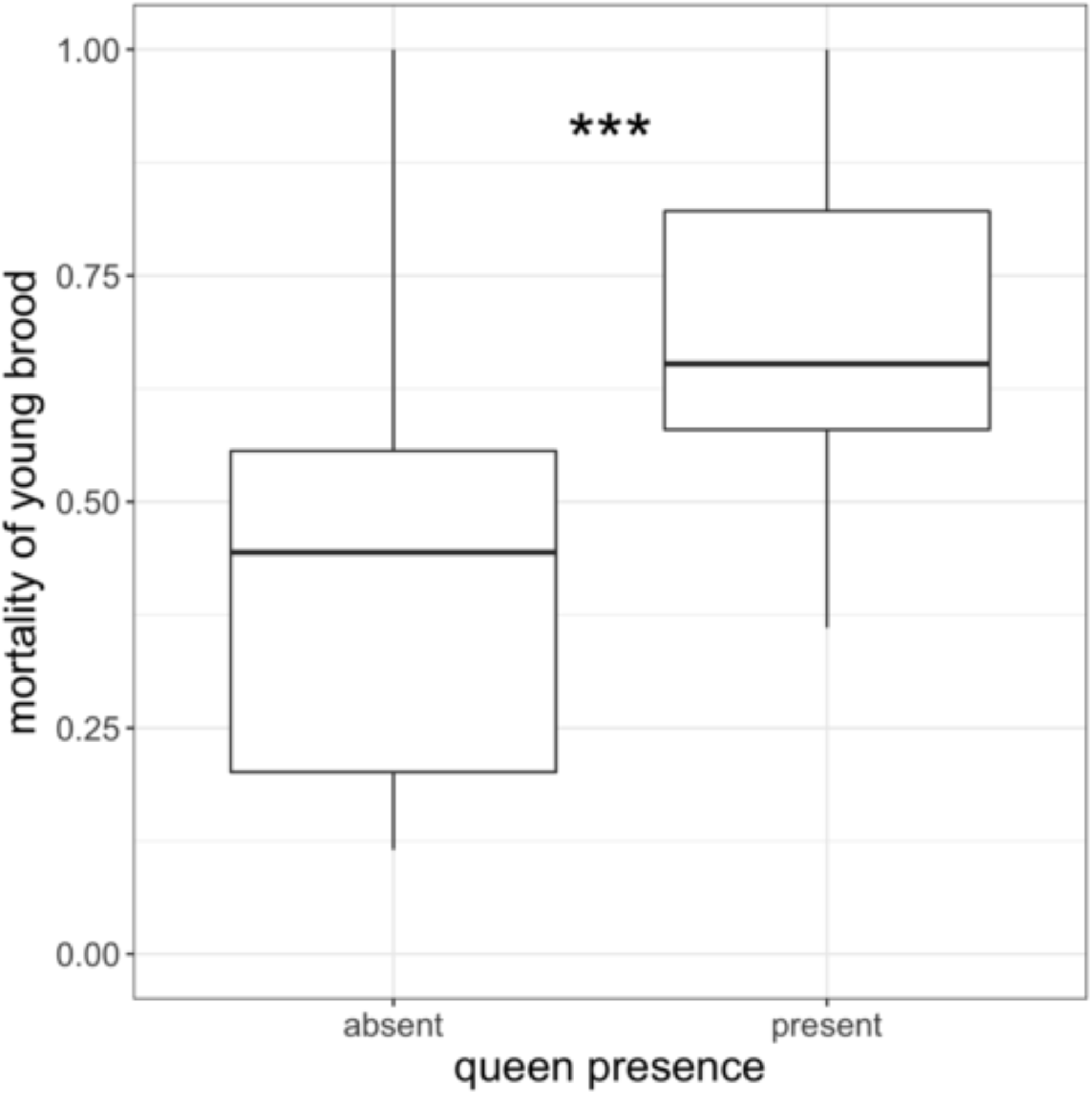
Young brood (eggs and 1st instar larvae) experienced higher mortality in colonies in which queens were re-introduced, indicating reproductive-destined young brood were culled. Equal numbers of age-matched (with 24 hrs.) eggs were reared in queenless colonies of equal size for one week, after which queens were replaced in half the colonies. In colonies where the queens were replaced (“present”), fewer eggs survived to the following week, at which point they had molted to the 2nd or 3rd larval instar. *** = p < 0.001 (LRT; binomial GLM)

## Discussion

The ability to efficiently allocate reproductive and somatic tasks to distinct members of colonies has allowed social insects to be ecologically diverse and dominant (Wilson 1971). Here, we find that colonies of the pharaoh ant *Monomorium pharaonis* dynamically alter resource allocation to fit colony needs in response to a combination of the numbers of workers and eggs contained in a colony. Thereby, colonies exhibit adaptive demography (Wilson 1985), making collective decisions about allocation based on observable present requirements for growth and fitness.

Colonies of *M. pharaonis* typically inhibit production of new reproductives (males and new queens, or gynes) in the presence of egg-laying queens (Edwards 1987; Edwards 1991; Schmidt et al. 2011). In our study, some colonies produced new reproductives despite the presence of queens, depending on the number of eggs contained in the colony two weeks before reproductive pupae were observed (Fig. 1, Fig. S1). Two weeks is about the amount of time it takes for an individual to grow from the 1st larval instar to the pupal stage (Peacock and Baxter 1950). Previously, we showed that caste is regulated by the end of the 1st larval instar, as larvae older than the 1st instar contained in colonies not producing reproductives were exclusively worker-destined (Warner et al. 2016). Therefore, data from our first experiment suggest that the caste of individuals that molt and continue growth after the 1st instar is dependent upon the number of eggs contained in the colony around the time of molting (i.e. when caste regulation occurs).

When colonies of *M. pharaonis* produce reproductives as a result of queen removal, they rear proportionally fewer reproductives with increasing colony size (Schmidt et al. 2011). In our first experiment, we found that rearing colonies with variable numbers of queens for 6 weeks (specifically 2,4, or 8 queens) was sufficient to cause a change in colony size (Fig. S2). This change in colony size subsequently impacted allocation to reproductives when queens were removed (Fig. S3). When we measured caste allocation in colonies while varying only one element of colony size at a time (number of workers, number of queens, or number of eggs), we found increasing egg number decreased allocation to reproductives (Fig. 2), while increasing the number of workers increased reproductive allocation (Fig. 3).

Altogether, our results suggest that the ratio of worker number to egg number regulates caste allocation. As the ratio of workers to eggs increases, colonies should attempt to produce more reproductives capable of rearing more eggs that then will act to reduce the worker to egg ratio. Alternatively, if there are too few workers to rear all available eggs, allocating resources to workers is optimal (Oster and Wilson 1978). Interestingly, reproductive pupae were observed later in colonies with more eggs (Fig. 4a) or fewer workers (Fig. 4b). Based on these results, colonies containing lower worker:egg ratios produced proportionally fewer males and gynes because they began rearing reproductives later (i.e. inhibition of male/gyne production was halted later or generally weaker).

It is unknown whether female caste (gyne versus worker) in *M. pharaonis* is determined in the egg stage, whether genetically (Anderson et al. 2008; Schwander et al. 2010) or blastogenically (i.e. via compounds endowed by the queen) (Bourke and Ratnieks 1999; Fersch et al. 2000; Schwander et al. 2008; Libbrecht et al. 2013), or whether it is determined nutritionally in the 1st larval instar (Warner et al. 2016) (note, however, that in the congeneric *M. emersoni*, female caste appears to be determined in the egg stage (Khila and Abouheif 2010)). In contrast, males, being haploid across hymenopterans, must be constantly produced and culled (Edwards 1987; Schmidt et al. 2011; Warner et al. 2016). Brood culling, particularly that of genetically-determined males, is common throughout ants (Aron et al. 1994, 1995; Passera and Aron 1996; Keller et al. 1996). If female caste is determined genetically or blastogenically (so eggs are strictly either gyne-, male-, or worker-destined), that would imply that workers cull excess gynes as well as males when egg density in colonies is high. Our data indicate workers indeed cull gyne-destined eggs, as larval mortality was about 40% greater in colonies in which queens were replaced compared to colonies that remained queenless (Fig. 5). This percentage approximates the overall allocation to reproductives in colonies that exhibited high reproductive ratios across our experiments.

Based on our empirical data, we have developed a simple conceptual model (Fig. 6). We propose that the likelihood of a given reproductive-destined 1st instar larva to pupate (rather than to be culled; yellow arrow) is a function of the ratio of worker number to egg number at that particular time (mediated pheremonally; blue arrow). This simple rule has important downstream consequences: 1) if the worker:egg ratio is low enough, no reproductive L1 are allowed to molt to L2 (culling of 100% in Fig. 6b). 2) If the worker:egg ratio increases in the presence of queens, colonies begin producing reproductives at a low rate (culling stimulation drops below 100%, allocation to reproductives increases from zero). 3) If egg production halts, the worker:egg ratio will gradually increase. The more eggs present (or the smaller the number of workers), the longer this increase takes, which in turn results in lower proportional allocation to reproductives with lower initial worker:egg ratios. This explains the reduced reproductive allocation in queenless colonies of increasing size (Schmidt et al. 2011), provided egg number increases faster than the number of workers. Indeed, this is likely the case in colonies not already producing reproductives, which is typically thought of as the default state of *M. pharaonis* colonies. Our model is also consistent with past results in *M. pharaonis* showing that colonies started with fewer queens produce new queens and males earlier (Tay et al. 2014) and that older worker-destined larvae (which in the current study varied concurrently with number of adult workers) enhance a colony’s ability to rear reproductives (Warner et al. 2016). Lastly, it is important to note that allocation decisions are still contingent upon nutrient availability; previously, we showed that colonies must have access to insect protein in order to produce males and gynes (Warner et al. 2016). Here, we have provided colonies with food *ad libitum* so that allocation decisions were made purely based on colony demography rather than nutritional constraints.

**Fig. 6.**
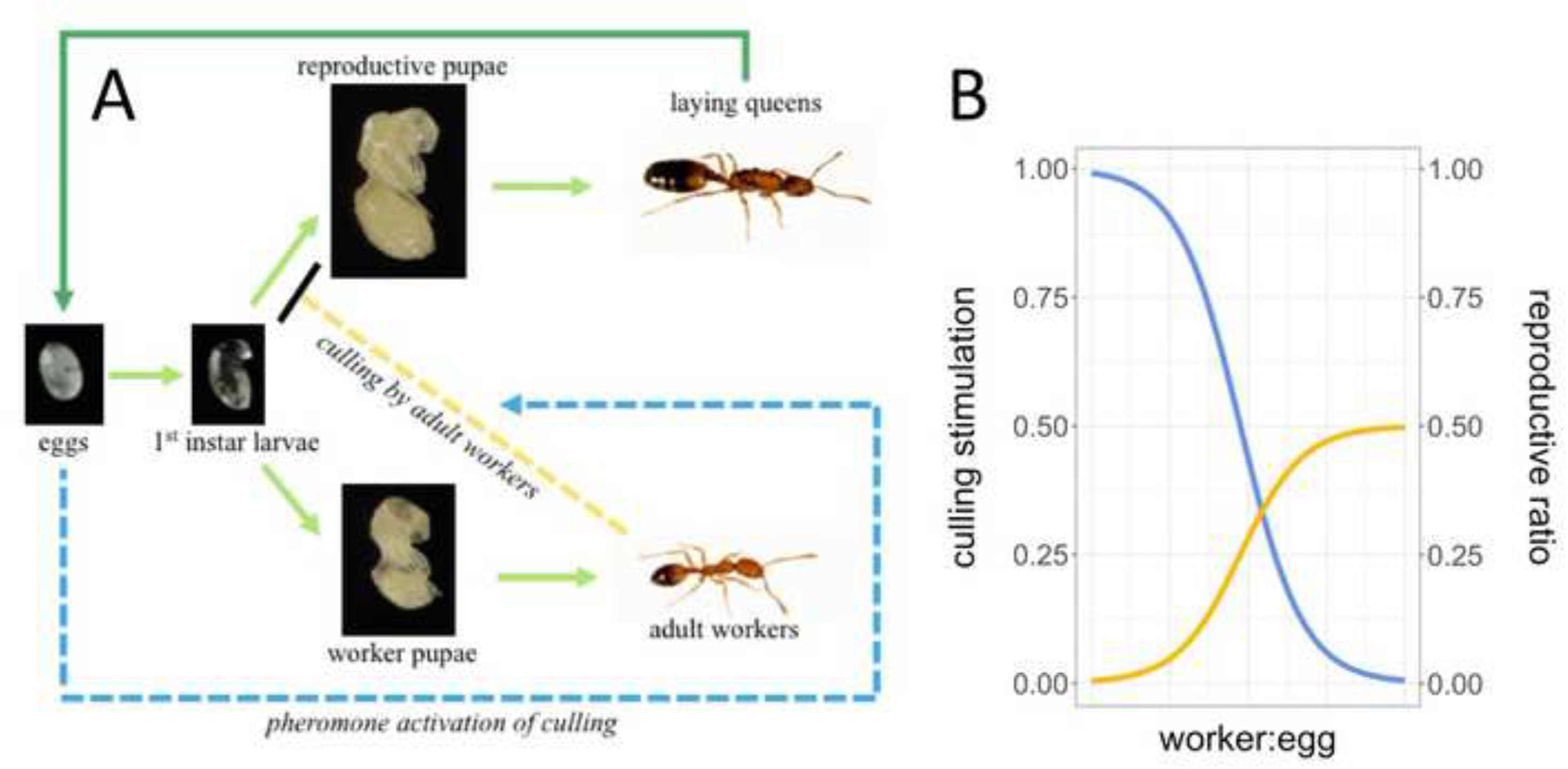
Conceptual model based on empirical results. A) Eggs promote (dotted blue arrow) worker culling (dotted yellow arrow) of reproductive-destined larvae before larvae reach the second instar. Solid arrows refer to development (light green) and egg laying (dark green), dashed arrows arrows refer to social interaction (culling; yellow) and communication (pheromonal stimulation of culling; blue). B) The stimulation of culling is tied to the proportion of pupae a colony produces that are males or new queens (reproductive ratio). Under this model, the reproductive ratio (solid blue line) is maximized (here, shown as 50%) and culling stimulation (solid yellow line) minimized at a high worker:egg ratio. At such a high worker:egg ratio, no culling occurs or culling is indiscriminate of larval caste. The maximum reproductive ratio is therefore equal to the proportion of queen-and male-destined eggs among all eggs laid by queens

The inhibition of gyne and male production in *M. pharaonis* is thought to be driven by the presence of freshly laid eggs, likely via a pheromone signal (Edwards 1987). In concordance with this theory, we found that the ratio of worker:eggs, rather than the number of queens, affects reproductive allocation, both qualitatively and quantitatively. This is consistent with the action of pheromonal inhibition, as a lower egg density within colonies dilutes the inhibitory power (analogous to reports of sporadic male production in larger argentine ant colonies (Passera et al. 1988)). Adult workers likely assess current colony demography based on social interactions with other adult workers as well as with eggs and larvae. Importantly, task allocation in insect societies is governed by rates and types of recent social encounters by individual colony members (Gordon 1996; Gordon and Mehdiabadi 1999). While each colony member can assess only a small amount of information about the colony state, integration of many members’ limited knowledge can result in optimal group behavior (Pacala et al. 1996), as exhibited in the present study.

Many social insect societies regulate production of reproductives (by both queens and workers) pheremonally. Queen pheromone signalling controls worker reproduction in many species (Holman et al. 2010, 2013; Van Oystaeyen et al. 2014) as well as the development of new queens in species that found colonies through budding (Vargo and Passera 1991, 1992; Boulay et al. 2007). While queen pheromone is often transmitted through direct contact with the queen (Vargo and Passera 1991, 1992; Holman et al. 2010, 2013; Van Oystaeyen et al. 2014), fertility-signalling compounds can also be present on eggs (Endler et al. 2004). Fertility signaling via compounds laid on eggs is functionally similar to the regulation of soldier production in *Pheidole* by pheromones produced by soldiers. In each case, the signal regulating allocation is proportional to the present demography of the colony, which reinforces adaptive demography (Wilson 1985). In this way, signaling appears to be honest, in agreement with the vast majority of fertility signaling in other social insect species (Holman et al. 2010, 2013; Holman 2012; Van Oystaeyen et al. 2014)

We found no evidence of a maternal effect on reproductive allocation that varies according to the number of queens present in colonies (Fig. S4). In some ants, queens provision diploid eggs with varying levels of compounds that bias caste (Anderson et al. 2008; Schwander et al. 2010). For example, queens in *Pogonomyrmex rugosus* that have overwintered provision eggs with varying levels of vitellogenin, which directly impacts caste determination (Libbrecht et al. 2013). We reasoned that the drop in reproductive allocation with increasing queen number (Fig. S3) could be a result of such a maternal effect. If queen-queen inhibition occurs, either through competition for nutrients or through pheromonal signaling (Vargo 1992), number of queens could be associated with a propensity to lay worker-biased eggs. However, reproductive allocation did not differ based on queen number when other factors were controlled for (Fig. S4). Notably, it is still possible that female caste is determined blastogenically, but that caste allocation is not plastic based on the environment queens experience (at least in the manner we have perturbed it).

Organisms commonly employ bet-hedging strategies, in which individuals sacrifice maximum growth rate in a single environment in order to survive and reproduce in under environmental fluctuation (Cohen 1966; Philippi and Seger 1989). The strategy presented here is clearly a form of bet-hedging. Because queens constantly produce eggs of all castes, colonies can react to shifts in egg density by utilizing present resources. However, because queens of colonies producing solely workers lay abundant eggs and individuals are culled extremely early (before substantial resources have been invested in them) and are likely eaten (Bourke 1991; Tschinkel 1993; Aron et al. 1994), the cost of bet-hedging in practice is low.

The bet-hedging strategy described here may be widely applied in other ant societies which form colonies through budding. Because queens do not disperse separately from the workforce, gyne production in budding societies is limited by local resource competition (among gynes for workers to rear their eggs and provide them with nutrition) (Pamilo 1991b; Bourke and Franks 1995; Pearcy and Aron 2006; Schmidt et al. 2011; Villalta et al. 2015). Accordingly, new gynes are reared only when the reproductive capacity of current queens diminishes (Vargo and Fletcher 1986, 1987; Arcila et al. 2002; Brown et al. 2002; Boulay et al. 2009; Schmidt et al. 2011), and gyne-destined larvae are culled when introduced to colonies containing laying queens (Edwards 1991; Villalta et al. 2016). This strategy can be thought as “adaptive demography” (Wilson 1985), particularly in the case of polygynous species, for which growth and reproduction are dependent upon numbers of both queens and workers (Pamilo 1991a; Cronin et al. 2013). This dependence necessitates the ability to rapidly shift female caste allocation in response to demographic disturbance. Polygynous budding ants include many widely successful invasive species that thrive in the presence of anthropogenic disturbance (Helanterä et al. 2009) and readily dominate habitats (Hölldobler and Wilson 1977; Boulay et al. 2014). Because colony propagules containing only workers and larvae may establish a new colony, bet-hedging (in the form of always containing young individuals that may develop into all castes) allows colonies to respond to and even thrive as a result of frequent disturbance.

It is reasonable to think that rapid social mechanisms promoting queen turnover are particularly important in polygynous species, as queens may senesce earlier, due to diminished selection for long lifespans (Keller and Genoud 1997). Indeed, older queens of *M. pharaonis* and *L. humile* lay lighter (presumably more poorly provisioned) eggs (Petersen-Braun 1977; Keller and Passera 1990) and polygynous ant queens in general exhibit much shorter lifespans (Hölldobler and Wilson 1977; Keller 1995; Keller and Genoud 1997). Problematically, if queens senesce slowly or asynchronously, the decline in egg number may be not be sharp enough for colonies to cross the worker:egg threshold to begin to produce new reproductives. This would result in a slow decline of worker number and colony collapse.

Other polygynous budding species display strategies that help avoid this fate. Because they engage in mating flights, colonies of the polygynous form of *Solenopsis invicta* experience a continual influx of new, re-accepted queens (Glancey and Lofgren 1988). Colonies of *L. humile* kill up to 90% of queens annually, before seasonal production of reproductives (Keller et al. 1989). This culling reduces the efficacy of pheromonal inhibition (Vargo and Passera 1991), allowing rearing of mass numbers of reproductives, replenishing the colony with young queens. Unlike *S. invita* and *L. humile*, *M. pharaonis* colonies are not known to cull adult queens and reproductives mate in the nest (Fowler et al. 1993), so colonies don’t display obvious mechanisms to prevent gradual senescence. Therefore, it is possible they are actually dependent upon disturbance to thrive, much in the way that plants (e.g. those that require fire to germinate) can be dependent on disturbance to reproduce (Howe and Smallwood 1982; Keeley 1987, 1991).

## Conclusion

Based on results from a series of manipulative experiments, we have devised a model describing the social regulation of caste allocation in *M. pharaonis*. We propose that workers cull queen-and male-destined larvae when egg density within colonies is sufficiently high, as signalled by pheromones on eggs. These newly-laid eggs are an accurate indicator of the presence of fertile queens. Colonies reduce or cease culling and begin to produce males and queens when egg density declines. Colonies vary in caste allocation as a function of the timing of the decline in egg density and culling. The social regulatory mechanisms presented here constitute a bet-hedging strategy that is likely active in other ant societies in which queens do not disperse. Colonies bet-hedge by retaining the ability to produce new queens and males but do so only when the fertility of existing queens is lowered. Because colonies regulate the caste fate of individual larvae early in larval development, colonies do not waste resources (e.g., investing in new reproductives when fertile queens already exist) and colonies can invest nearly all resources in colony growth while retaining flexibility to shift to investment in reproduction whenever necessary.

## Acknowledgements

We thank Rohini Singh and Justin Walsh for comments on previous drafts of the manuscript. This research was funded by National Science Foundation award IOS-1452520. MRW was funded by National Science Foundation award DGE-1321851.

---

**Fig. S1** Colonies that were established with similar sizes (~1.2 mL; ~175 adult workers) but different numbers of queens differed in caste allocation. A) Colonies began producing reproductives before queen removal when started with fewer queens. B) Colonies which we observed producing reproductive pupae in the presence of queens contained fewer eggs two weeks prior to the given observation (approximately the developmental time between caste regulation of 1st instar larvae and pupation), demonstrating that number of eggs affects caste regulation

**Fig. S2** Initially equal-sized (~1.2 mL; ~175 adult workers) colonies differed in colony productivity according to the number of queens they contained at colony creation. At the time of queen removal (6 weeks after colony creation), there was a positive relationship between the number of queens and the number of A) worker pupae, and B) eggs present in colonies, demonstrating that queen number strongly affects colony productivity

**Fig. S3** When queens were removed, caste allocation differed between experimental colonies created with two versus four or eight queens. Colonies created with two queens produced higher A) caste ratios (gynes/[gynes+workers]) and B) reproductive ratios ((gynes+males)/(gynes+males+workers)) than those created with four or eight queens. This was the result of increased production of worker pupae in colonies with four and eight queens (B)

**Fig. S4** When equal-sized colonies were established with two or eight queens, the number of queens that laid the eggs did not affect production of any caste after queen removal. After removing queens, the number of young brood (eggs and 1st instar larvae) was standardized, such that colonies were demographically equivalent, but differed in the number of queens that laid eggs. This demonstrates that the the shift in caste allocation accompanying increased colony size is not dependent upon a maternal effect

